# TransStop, a genomic language model for the pan-drug prediction of translational readthrough efficacy

**DOI:** 10.1101/2025.08.29.672857

**Authors:** Nicolas Haas, Arnaud Kress, Julie D. Thompson, Olivier Poch

## Abstract

**Motivation:** Premature termination codons (PTCs) are a major cause of genetic diseases, but the efficacy of therapeutic readthrough agents is highly context-dependent. While linear models have shown promise in predicting readthrough efficiency, they may not fully capture the complex, non-linear interactions between sequence context and drug activity.

**Methods:** We developed TransStop, a transformer-based pan-drug model, trained on a dataset of ∼5,400 PTCs and eight readthrough compounds. The model learns sequence representations and incorporates learnable embeddings to capture drug-specific effects, allowing a single model to predict efficacy for multiple drugs.

**Results:** Our model achieved a global R^2^=0.94 on a held-out test set. Visualizations of the learned embeddings revealed a deep understanding of biological principles, including the distinct clustering of stop codon types and grouping of drugs by mechanism of action. *In silico* saturation mutagenesis and epistasis analyses uncovered complex, non-additive sequence determinants of readthrough. We generated 32.7 million predictions across the human genome, covering all possible PTCs. Analysis of these genome-wide predictions revealed strong drug specializations for specific stop codon contexts and identified key areas of disagreement with previous models, particularly for UGA codons, where our model predicts a more effective drug in thousands of cases. The TransStop model represents a significant advancement in the prediction of translational readthrough efficiency. Its superior accuracy and the biological insights derived from its applications provide a powerful tool for guiding clinical trial design, drug development, and personalized patient treatment.

**Availability and Implementation:** Source code: https://github.com/Dichopsis/TransStop. Model: https://huggingface.co/Dichopsis/TransStop.Genome-wide predictions: https://doi.org/10.5281/zenodo.16918476.

## Introduction

Premature termination codons (PTCs) are responsible for approximately 12% of rare human genetic diseases (Mort et al. 2008) and have been observed in ∼10% of tumor suppressor genes, including the *TP53* gene that is involved in more than 50% of human cancers (Bidou et al. 2017). Mutations leading to PTCs result in the synthesis of truncated, often non-functional or dominant-negative proteins, and can trigger nonsense-mediated mRNA decay (NMD), further reducing the potential levels of protein production (Lejeune* 2023, Kurosaki, Popp and Maquat 2019, Karousis, Nasif and Mühlemann 2016). A promising therapeutic strategy to counteract the effect of PTCs is to promote translational readthrough, a process in which the ribosome bypasses the PTC, allowing for the synthesis of a full-length, functional protein (Dabrowski, Bukowy-Bieryllo and Zietkiewicz 2018, Morais, Adachi and Yu 2020).

The efficacy of readthrough-inducing small molecules is highly dependent on the PTC’s type (UAA, UAG or UGA), local sequence context (Wangen and Green 2020) and the specific drug used (Trzaska et al. 2020, Bidou et al. 2022). Recently, (Toledano, Supek and Lehner 2024) performed a landmark experimental study, quantifying the readthrough of approximately 5,800 pathogenic human PTCs for eight different drugs. Their work revealed that different drugs promote the readthrough of distinct and complementary subsets of PTCs. Using this extensive dataset, they developed an interpretable logistic regression model that successfully predicted drug-induced readthrough, achieving a high correlation (R^2^ = 0.83) for a pan-drug model.

While groundbreaking, the predictive models from Toledano et al. were based on logistic regression, which may not fully capture the complex, non-linear interactions between the sequence context and the drug’s mechanism of action. To address this, we developed TransStop, a genomic Language Model (gLM) based on the transformer architecture (Vaswani et al. 2023, Boshar et al. 2024, Consens et al. 2025). Our model was fine-tuned from the publicly available, pre-trained “nucleotide-transformer-v2-500m-multi-species” model (Dalla-Torre et al. 2025), allowing us to leverage its powerful, pre-existing knowledge of DNA sequence patterns.

In this study, we present a novel, pan-drug predictive model based on a nucleotide transformer architecture. Our primary goals were to: 1) create a single, unified model capable of predicting readthrough efficiency for multiple drugs simultaneously by learning drug-specific embeddings; 2) leverage the deep learning capabilities of a transformer to capture complex sequence-drug interactions that linear models may miss; 3) rigorously compare the performance of our transformer-based model against the state-of-the-art linear models on a held-out test set; and 4) deploy our model for genome-wide predictions to identify novel therapeutic opportunities and illustrate its use with a case study on the cystic fibrosis transmembrane conductance regulator (CFTR) gene.

## System and Methods

### Data acquisition and preprocessing

The primary dataset for this study was sourced from Toledano et al. and contains readthrough efficiency measurements for ∼5,800 PTCs across eight drugs and an untreated control. The PTCs were originally extracted from the ClinVar database (Landrum et al. 2014) and were reliably annotated as pathogenic. The eight selected drugs covered different classes of small molecules: SRI, SJ6986 and CC90009 are eRF1/eRF3 inhibitors (Lee et al. 2022, Sharma et al. 2021, Surka et al. 2021); DAP interferes with the activity of FTSJ1(tRNA-specific 2’-O-methyltransferase) (Trzaska et al. 2020); G418 and gentamicin are aminoglycosides that bind to the decoding center of the ribosome (Howard, Frizzell and Bedwell 1996); Clitocine (Friesen et al. 2017) and FUr (Palomar-Siles et al. 2022) are nucleoside analogs that are incorporated into the mRNA. The raw data was downloaded from the associated Figshare repository (https://figshare.com/articles/dataset/Other_objects/25138712).

The initial dataset for each drug was filtered using the same criteria as Toledano et al., ensuring comparability between predictive models. Specifically, we retained only measurements from a single replicate with read counts exceeding 15, excluded viral entries, and removed records lacking readthrough values, yielding approximately 5,400 PTCs per drug.

All drug-specific data were then integrated and concatenated into a single dataset. A critical feature engineering step was performed to create sequence contexts of varying lengths around the central stop codon. The target readthrough variable was log(x+1)- transformed to stabilize variance and normalize the distribution for model training. Finally, the dataset was split into training (80%), validation (10%), and test (10%) sets. This split was stratified by drug to ensure that the distribution of compounds was consistent across all datasets.

## Model architecture

Our predictive model, TransStop, is based on a deep learning architecture that integrates genomic sequence and drug information to predict readthrough efficiency. The model’s foundation is the pre-trained `InstaDeepAI/nucleotide-transformer-v2-500m-multi-species` transformer (Dalla-Torre et al. 2025), which acts as a powerful sequence feature extractor.

A key innovation in TransStop is the use of a cross-attention mechanism to dynamically model the interaction between the drug and the nucleotide sequence. The architecture comprises three core components:

1. **Sequence Encoder**: The base nucleotide transformer processes the input RNA sequence (e.g., ±6 nucleotides around the PTC) and generates a sequence of contextual embeddings, one for each nucleotide token.
2. **Drug Embedding**: A learnable embedding layer converts the identity of a given drug into a dense vector representation. This drug embedding is learned during training and captures the specific properties of each compound.
3. **Cross-Attention Regression Head**: Instead of simple concatenation, a custom regression head utilizes a cross-attention layer. In this mechanism, the drug embedding serves as the “query” vector. The query attends to the sequence of nucleotide embeddings (the “keys” and “values”), allowing the model to dynamically weigh the importance of different parts of the sequence context for a specific drug. The output of the cross-attention layer, a drug-conditioned sequence representation, is then passed through a feed-forward neural network (two linear layers with a ReLU activation and dropout) to produce the final scalar prediction of readthrough efficiency.

### Model training and hyperparameter optimization

The model development process involved two main stages: an ablation study to find the optimal sequence context length and a hyperparameter optimization to fine-tune the model configuration.

First, the ablation study was performed to determine the most informative sequence context length. Different models were trained using sequence contexts of varying lengths (±0, ±3, ±6, ±9, ±21, and ±72 nucleotides around the stop codon). The primary metric for this evaluation was the coefficient of determination (R^2^) on the validation set, provided in Table 1. We selected the shortest context where the R^2^ was less than 0.01 lower than that of the context with the highest R^2^. This resulted in the selection of the ±6-nucleotide context (R^2^=0.927) for the final model.

**Table 1.** Coefficients of determination (R^2^) for different sequence context lengths obtained on the validation data set.

| Context length<br>(nucleotides) | $\pm 0$ | $\pm 3$ | $\pm 6$ | $\pm 9$ | $\pm 21$ | $\pm 72$ |
| --- | --- | --- | --- | --- | --- | --- |
| $R^2$ | 0.613 | 0.899 | 0.927 | 0.928 | 0.930 | 0.930 |

Next, we conducted a comprehensive hyperparameter search using the Optuna framework (Akiba et al. 2019). The optimization process, run for 30 trials, aimed to maximize the R^2^ score on the validation set. The search identified the following optimal hyperparameters: a learning rate of 2.81e-05, a batch size of 16, weight decay of 0.071, a warm-up ratio of 0.12, and a cosine learning rate scheduler. For the regression head architecture, the optimal configuration was a hidden layer size of 512, a drug embedding size of 32, and a dropout rate of 0.17.

After identifying the best hyperparameter set, a final production model was trained on the combined training and validation datasets to maximize the data available for learning. The final model, along with its associated tokenizer and drug-to-integer mapping, was used for downstream inference and analysis.

### Genome-wide inference and analysis

To demonstrate the model’s utility for large-scale discovery, we performed genome-wide inference on all possible PTCs in the human genome. The input data was sourced from Toledano et al., and contained all potential single-, bi- or tri-nucleotide variants resulting in an amino acid change to a PTC. This process involved extracting the appropriate sequence context (±6 nucleotides) for each potential PTC and then employing a multi-GPU pipeline to accelerate the prediction process for all eight drugs. The resulting 32.7 million predictions were merged with the original genomic data and saved as a comprehensive file for analysis.

## Results

### Development of a pan-drug transformer model to predict stop codon readthrough efficiency

We developed a pan-drug, transformer-based model, TransStop, to predict the efficiency of translational readthrough. The model consists of a sequence encoder to process the RNA sequence context (±6 nucleotides) around a PTC, and a learnable embedding layer that converts the identity of a given drug into a dense vector representation, capturing the specific properties of the compound. A drug-sequence cross-attention mechanism enables the model to learn not just what a sequence looks like, but how a particular drug “sees” that sequence, providing a more nuanced and more powerful framework for predicting drug-specific efficiency. The model was trained and optimized on a dataset containing experimental readthrough efficiency measurements for ∼5,400 pathogenic PTCs and eight drugs.

### Evaluation of the accuracy of the TransStop model and comparison with baseline models

The TransStop model was evaluated on a held-out test set, and demonstrated remarkable overall performance, achieving a coefficient of determination (R^2^) of 0.94 (Fig. 1A). This high correlation confirms that our model reliably predicts the scale of the therapeutic effect.

**Figure 1.**
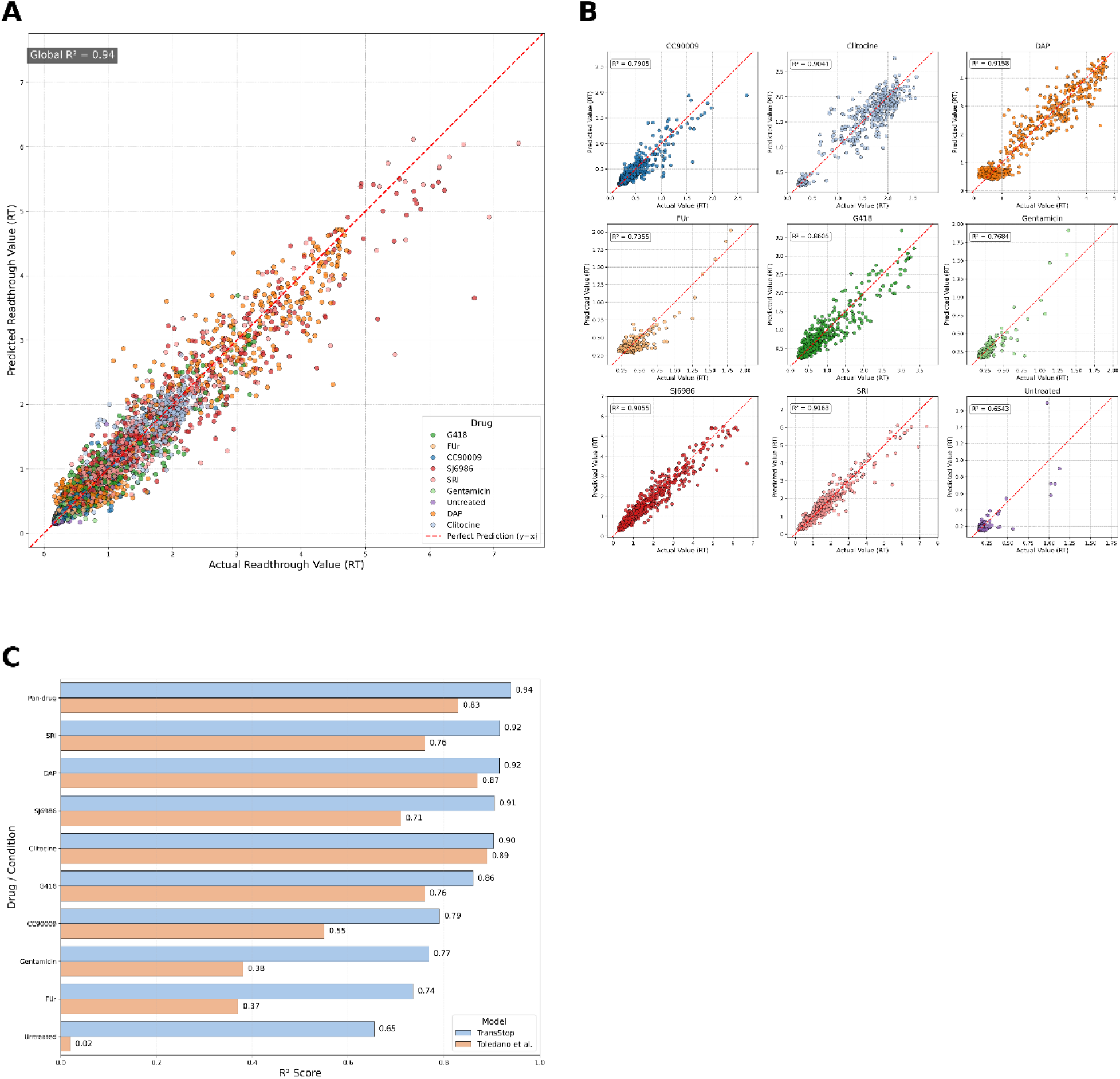
Prediction accuracy of the pan-drug transformer model TransStop and comparison with baseline models. (A) Global scatter plot showing the correlation between model-predicted readthrough (RT) values and experimentally measured actual values from the test set. Each point represents a unique sequence-drug combination. The dashed red line (y=x) indicates a perfect prediction. The global coefficient of determination (R^2^) is displayed in the top-left corner. (B) Scatter plots showing the predictive performance for each individual drug, as well as for the “Untreated” condition. The R^2^ is calculated and displayed for each condition. (C) Bar chart of performance (R^2^ scores) for the TransStop model (blue) and the logistic regression model from Toledano et al. (orange) for each drug and for the overall “Pan-drug” performance. Data for our model is derived from the evaluation on the held-out test set.

At the individual drug level, the model showed robust individual performances (Fig. 1B). It achieved an accuracy (R^2^ > 0.90) for key drugs, including the eRF1/eRF3 inhibitors SRI (R^2^=0.92) and SJ6986 (R^2^=0.91), as well as for DAP (R^2^=0.92) and Clitocine (R^2^=0.90). This suggests the model is particularly adept at capturing the sequence-dependent mechanisms of these compounds. Performance remained strong for aminoglycosides like G418 (R^2^=0.86)/Gentamicin (R^2^=0.77) and other agents such as the eRF3 inhibitor CC90009 (R^2^=0.79).

Crucially, TransStop systematically outperformed the state-of-the-art logistic regression model from Toledano et al. across every tested condition (Fig. 1C). The overall pan-drug R^2^ of 0.94 represents a significant improvement over the baseline model with R^2^=0.83. The improvement in performance was most pronounced for targeted eRF1/eRF3 inhibitors like SJ6986 (0.91 vs. 0.71) and CC90009 (0.79 vs. 0.55), highlighting our model’s superior ability to capture the complex, non-linear sequence-drug interactions that govern the efficiency of modern therapeutic compounds.

### Interpretation of the TransStop model’s latent space in terms of biological features

To investigate the biological knowledge learned by TransStop, we visualized the drug-conditioned context vectors from the model’s latent space using Uniform Manifold Approximation and Projection (UMAP) (McInnes, Healy and Melville 2020). This analysis revealed that the model internally organizes the data based on fundamental biological and mechanistic principles in a completely unsupervised manner.

The most striking finding is that the model spontaneously partitions the sequence space by stop codon identity. When the UMAP projection is colored by stop codon type, three distinct clusters emerge, corresponding to UAA, UAG, and UGA (Fig. 2A). This emergent property demonstrates that the model identified the stop codon as the most salient feature for predicting readthrough efficiency. Furthermore, this organization is functionally relevant: coloring the same projection by readthrough values reveals clear gradients, with high-readthrough sequences occupying specific niches at the extremities of the codon clusters (Fig. 2B).

**Figure 2.**
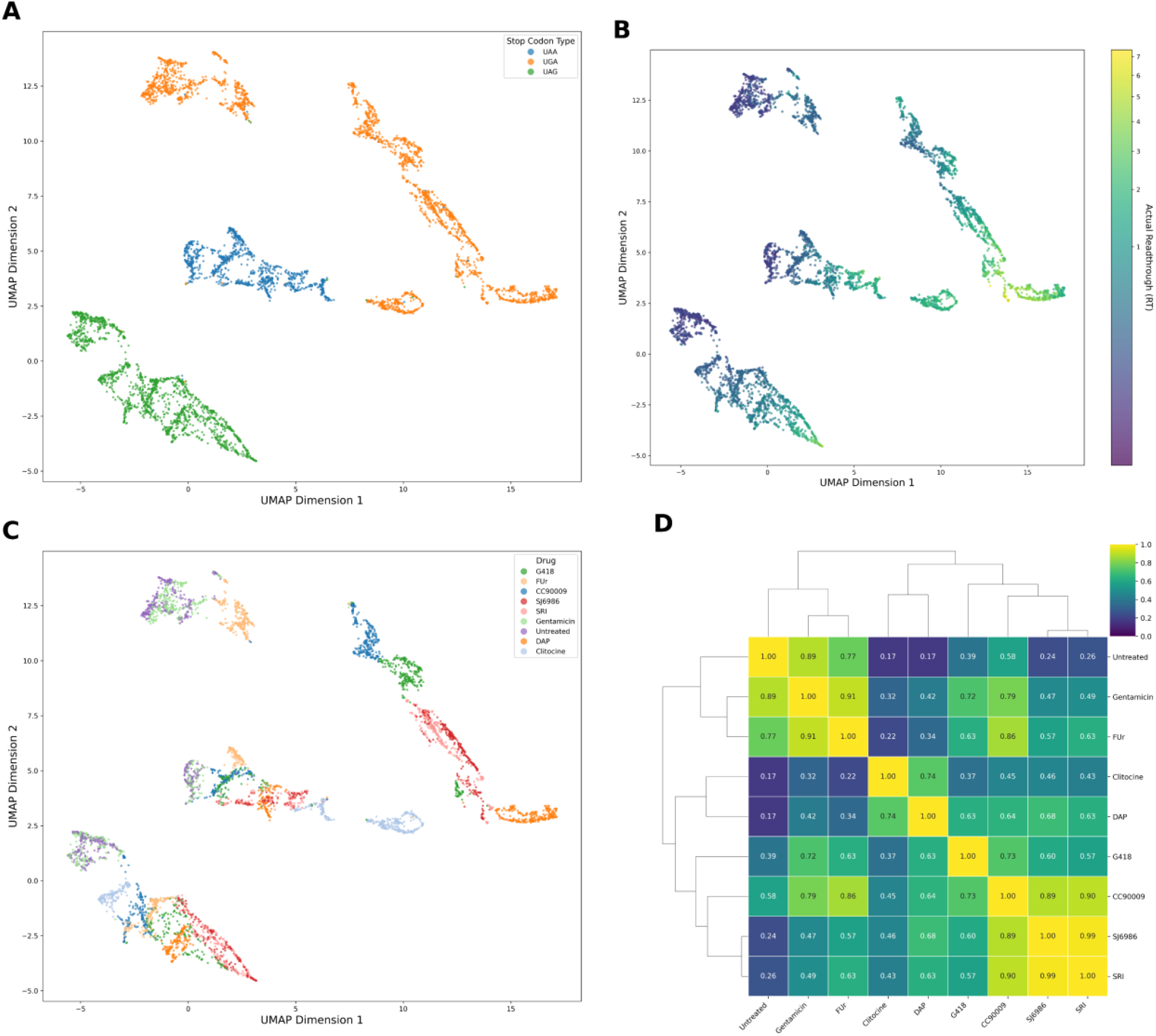
Biological interpretation of TransStop’s latent space. (A) The UMAP projection colored by the stop codon type (UAA, UAG, UGA) of the sequence. The plot reveals three distinct and well-separated clusters, corresponding to the three stop codon types. (B) The same UMAP projection, colored by the experimentally measured actual readthrough (RT) value. A color gradient from low (dark purple) to high (bright yellow) RT is visible, particularly at the extremities of the clusters. (C) The UMAP projection colored by drug shows how the representations for different drugs are organized within the primary stop codon clusters. (D) A hierarchical clustering (clustermap) of the drugs based on the functional similarity of their prediction profiles. The similarity was calculated as the Pearson correlation between the *in silico* readthrough predictions for each drug across all unique sequences in the test set.

The cross-attention mechanism allows the model to create drug-specific representations (Fig. 2C). While the global structure is predominated by the stop codon, the drug coloring shows that compounds form distinct, partially overlapping sub-patterns. This demonstrates that the model learns a universal representation of a sequence’s readthrough potential and then optimizes this representation based on the specific drug, capturing the unique drug-sequence interaction.

To quantify these relationships, we computed a functional similarity matrix by correlating the complete *in silico* readthrough profiles for all pairs of drugs (Fig. 2D). The resulting hierarchical clustering reflects known drug mechanisms. The eRF1/eRF3 inhibitors (SRI, SJ6986, and CC90009) form a distinct cluster with high inter-correlations (r > 0.89), indicating that the model has learned their shared functional signature. While the two aminoglycosides, G418 and Gentamicin, also show functional similarity as expected (r=0.72), the clustering reveals an even stronger functional link between Gentamicin and the nucleotide analog FUr (r=0.91). This suggests that our model has captured subtle convergences in their functional impact that go beyond simple mechanistic classification.

### Case study: *in silico* mutagenesis for DAP and a UGA Codon

To demonstrate the applicability of our model as an *in silico* tool for mechanistic investigation, we examined the sequence–function relationship in a specific therapeutic context: the action of the drug DAP on a UGA stop codon. We performed saturation mutagenesis on the highest-performing UGA-containing sequence for DAP in the test set, predicting the effect of every possible single nucleotide substitution (Fig. 3A). The analysis identified the nucleotide at the +4 position (immediately following the UGA codon) as a critical determinant. The reference ‘C’ at this position is optimal, and any mutation away from it is predicted to be highly detrimental to readthrough, a finding that aligns with experimental data identifying the +4 C as a key readthrough enhancer.

**Figure 3.**
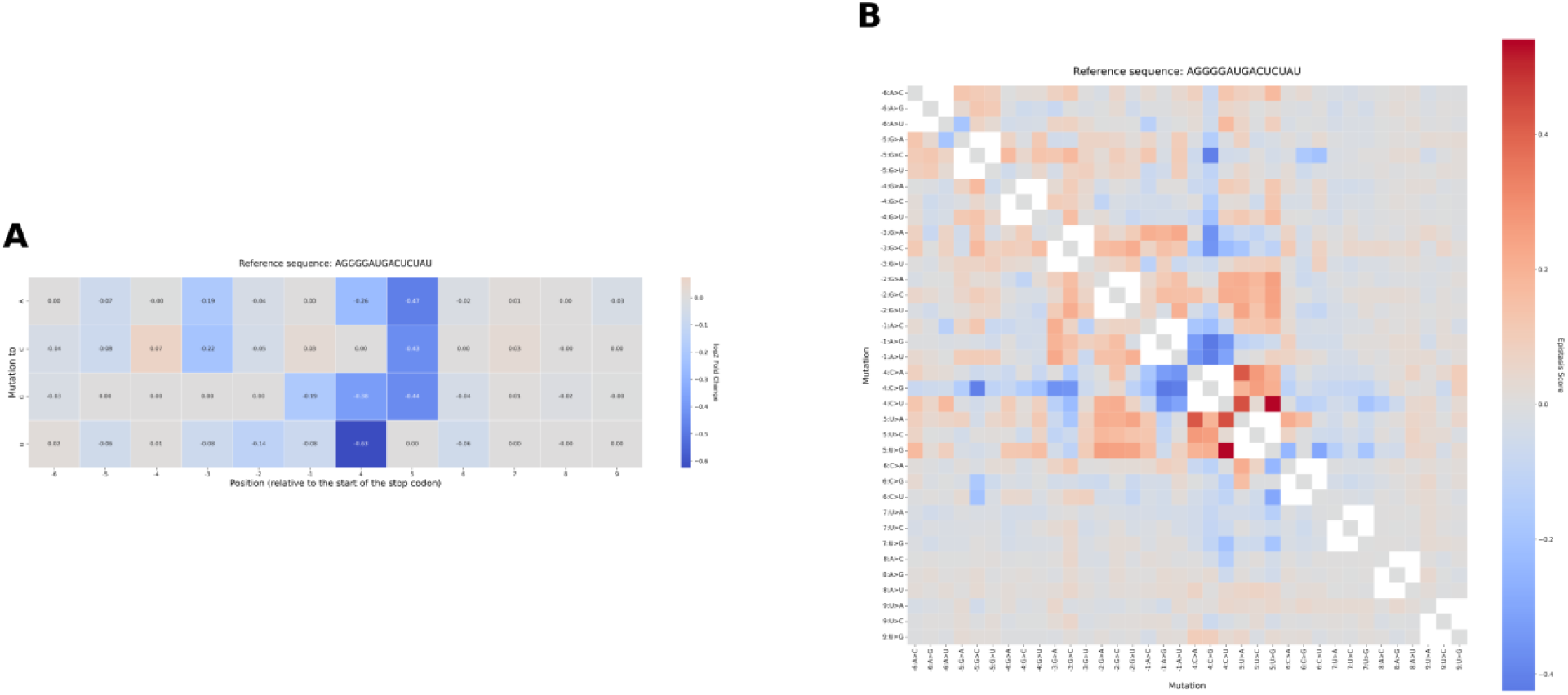
I*n silico* mutagenesis for DAP with a UGA PTC reveals key positional determinants and epistatic interactions. (A) A saturation mutagenesis heatmap showing the predicted impact of every possible single nucleotide substitution on readthrough efficiency. The effect is reported as the log2 fold change relative to the wild-type reference sequence with DAP treatment. Positions are numbered relative to the start of the UGA codon. Red cells indicate mutations that increase readthrough, while blue cells indicate those that decrease it. (B) An epistasis heatmap quantifying the non-additive effects of all possible double mutations in the context of DAP treatment. The epistasis score is calculated as the difference between the observed log2 fold change of the double mutant and the expected change from the sum of its individual constituent mutations. Red cells denote positive epistasis (synergy), while blue cells denote negative epistasis (antagonism).

To uncover higher-order sequence grammar, we then performed an epistasis analysis, predicting the effect of all possible double mutations (Fig. 3B). This analysis revealed a complex landscape of non-linear interactions specific to the DAP/UGA context. We identified synergistic interactions (positive epistasis, red squares), where combinations of mutations enhance readthrough more than the sum of their individual effects, and antagonistic interactions (negative epistasis, blue squares), where mutations clash to produce a suboptimal context. We thus infer that the transformer has captured non-additive rules, providing a deeper, mechanistic understanding of drug-sequence specificity that can inform the rational design of therapeutic contexts.

### Genome-Wide Prediction of readthrough levels using TransStop

Having validated the TransStop model’s performance and interpretability, we used it for genome-wide inference to predict the readthrough level of 32.7 million potential PTCs across the human genome. This large-scale analysis provides a comprehensive view of the therapeutic landscape, revealing specific drug specializations.

An analysis of the predicted best drug for each PTC confirms that therapeutic efficiency is highly conditional on the stop codon type (Fig. 4A). Three drugs, SJ6986 (33.6%), DAP (31.7%), and Clitocine (30.6%) dominate the landscape, together accounting for over 95% of the optimal predictions. Their dominance may be due to their specialization: Clitocine is the best predicted drug for the majority of UAA codons (accounting for 96.4% of best cases), DAP is the best predicted drug for most UGA codons (87.4%), and SJ6986 is the best predicted drug for most UAG codons (84.1%).

**Figure 4.**
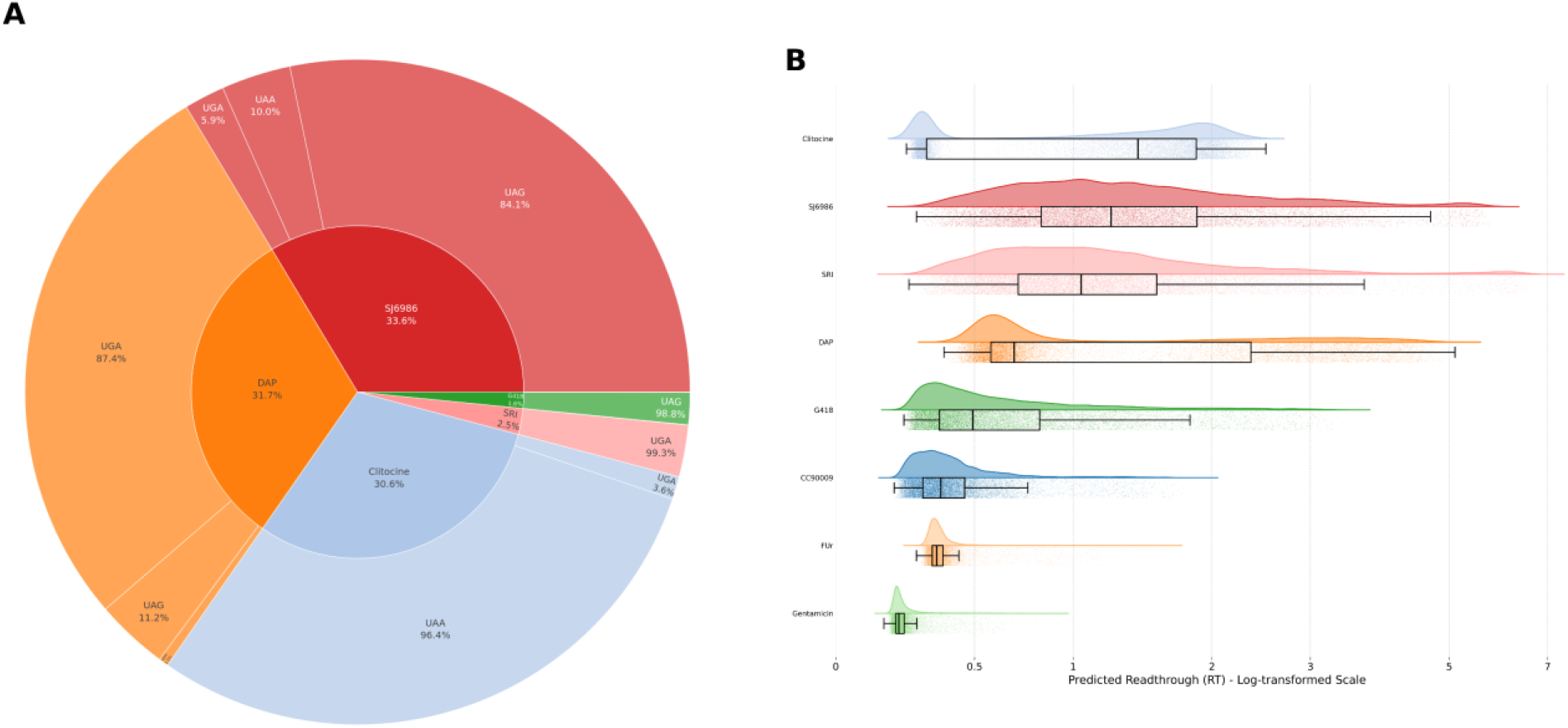
Analysis of genome-wide readthrough predictions. (A) An inverted sunburst plot visualizing the specialization of each drug for different stop codon types across all predicted PTCs in the human genome. The inner circle shows the proportion of all PTCs for which each drug is predicted to be the most effective. The outer ring details the breakdown of these PTCs by stop codon type (UAA, UAG, or UGA) for each drug, showing the percentage of the drug’s “best” cases that belong to each codon type. (B) A custom raincloud plot characterizing the distribution of predicted readthrough (RT) values for each drug. For each drug, the plot displays the probability density (the “cloud” or half-violin), a random sample of 100,000 individual data points to visualize density (the “rain” or strip plot), and a summary boxplot showing the median, interquartile range, and outlier whiskers. The x-axis is log-transformed to better visualize the distribution across different orders of magnitude.

To characterize the nature of each drug’s activity, we analyzed the distribution of their predicted readthrough values (Fig. 4B). The results distinguish between “high-potency specialists” and “broad-spectrum generalists.” Drugs like SJ6986, SRI, and DAP show heavy-tailed distributions, indicating that while they are effective for a narrow range of contexts, they can induce very high readthrough in those niches. In contrast, Clitocine exhibits a broader distribution with a higher median, suggesting it acts as a reliable generalist, inducing modest but consistent readthrough across a wide range of its preferred UAA contexts. Other drugs, such as the aminoglycosides, show efficacy profiles confined to a smaller number of PTCs, highlighting their more limited or niche applications.

### Comparison of the best predicted drug using TransStop and baseline models

To investigate whether our model provides novel, actionable insights, we compared its genome-wide predictions for the best therapeutic drug with those of the Toledano et al. model. While both models show high concordance for top specialist drugs like Clitocine, SJ6986 and DAP, the analysis revealed disagreements that highlight the advanced capabilities of our architecture (Fig. 5A).

**Figure 5.**
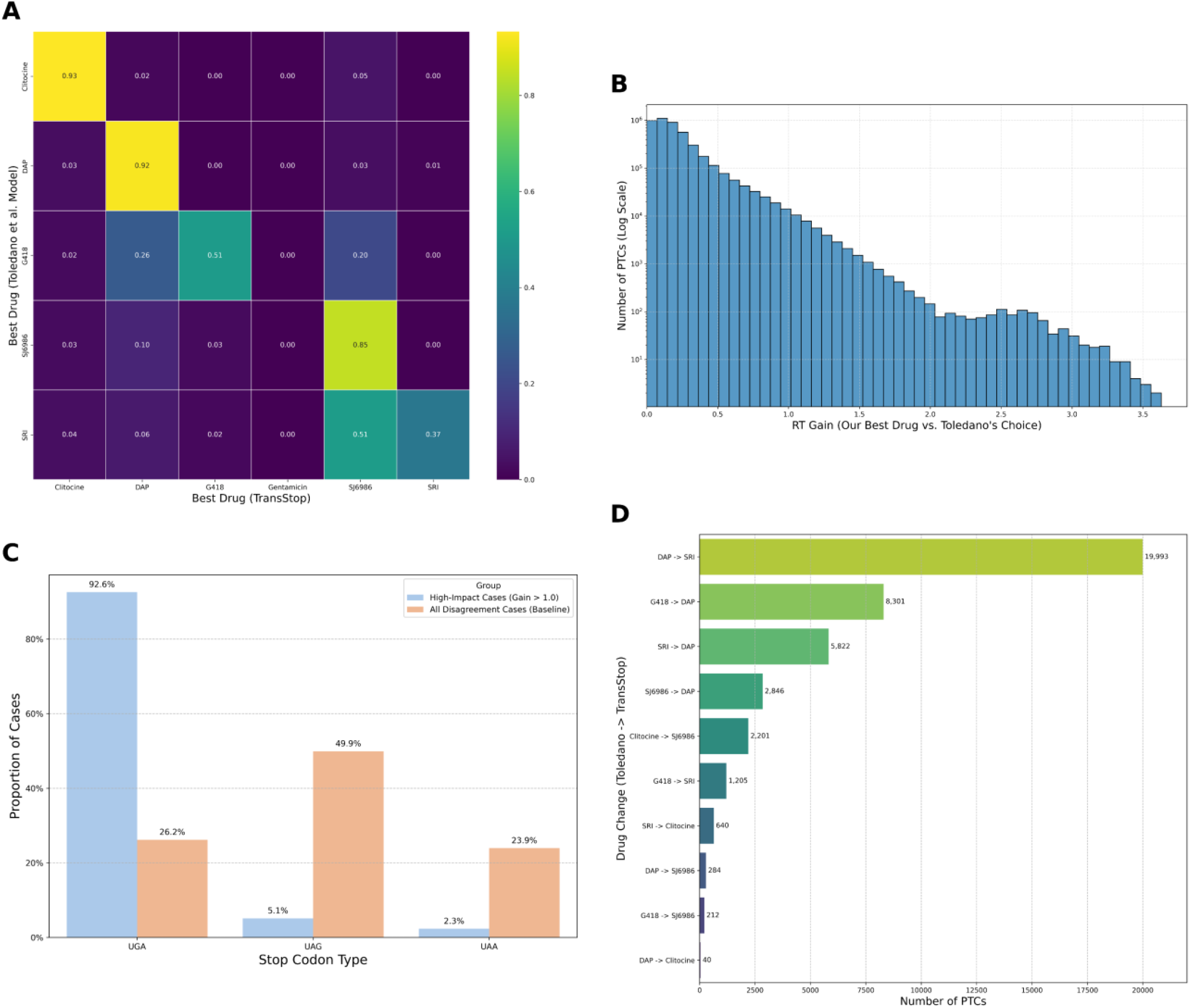
Comparison of therapeutic recommendations by TransStop and baseline models. (A) A confusion matrix showing the concordance in the prediction of the drug achieving the highest readthrough (RT). Each cell (i, j) indicates the proportion of cases where Toledano et al. predicted drug i as best compound and TransStop predicted drug j. The strong diagonal indicates general agreement for the top specialist drugs, but notable off-diagonal values highlight systematic disagreements. (B) A log-scale histogram of the predicted readthrough (RT) gain achieved by TransStop in all cases of disagreement. The x-axis represents the performance improvement (TransStop Best RT – Toledano et al. Best RT), showing that a large number of disagreements result in a positive gain. (C) A bar chart comparing the stop codon type distribution in high-impact disagreement cases (where TransStop’s predicted RT gain is > 1.0) versus the baseline distribution across all disagreement cases. (D) A bar chart of the top 10 most frequent drug switches in high-impact disagreement cases, indicating which therapeutic recommendations are most commonly revised by our model.

We quantified the “RT gain”, the predicted readthrough of our model’s best drug minus that of the Toledano model’s choice. The distribution of this gain shows 41,664 cases where a clinically meaningful improvement is observed (RT gain > 1.0), indicating that reliance on the previous model could lead to suboptimal therapeutic strategies in numerous instances (Fig. 5B).

A deeper analysis revealed that these high-impact disagreements are not random but are overwhelmingly driven by a specific biological context: the UGA stop codon. While UGA codons account for only 26.2% of all disagreements, they are responsible for 92.6% of the high-impact cases where our model predicts a substantial therapeutic gain (Fig. 5C). This suggests that our transformer-based model has a more sophisticated understanding of the complex sequence determinants surrounding UGA codons. The most common high-impact “drug switches” involve revisions from various drugs to the UGA-specialist DAP or the potent UAG/UGA-active drug SRI, providing concrete, testable hypotheses for improved therapeutic interventions (Fig. 5D).

### Predicted Therapeutic Profiles for Key CFTR Nonsense Mutations

As a final demonstration of its practical utility, we used TransStop to generate a “therapeutic matrix” for four clinically relevant nonsense mutation positions in the Cystic Fibrosis Transmembrane Conductance Regulator (CFTR) gene (Fig. 6). For the key nonsense mutations (i.e., G542X, R553X, R1162X, W1282X), we predicted the efficiency of all drugs for each of the three possible stop codons that could arise.

**Figure 6.**
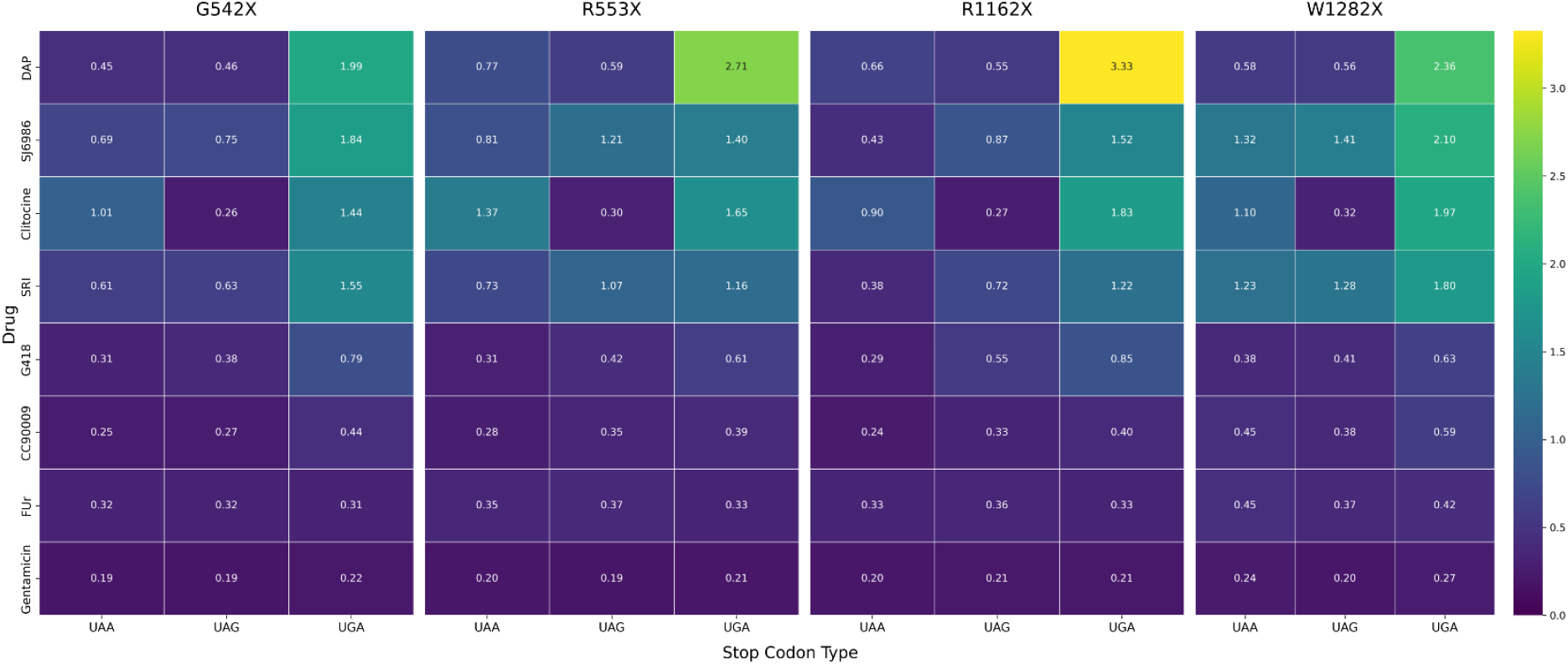
Predicted therapeutic profiles for key CFTR nonsense mutations. The four heatmaps correspond to the clinically relevant nonsense mutation positions in the CFTR gene (G542X, R553X, R1162X, and W1282X). For each position, the predicted readthrough (RT) efficiency for all tested drugs is shown (y-axis) for each of the three possible stop codon types (x-axis) that could result from a mutation at that site. Cell values and color intensity represent the predicted RT value, providing a direct “therapeutic matrix” for each mutation location.

The results indicate that efficiency is highly dependent on the complete mutational context (genomic position plus stop codon type). For example, a UGA codon at position R1162X is predicted to be exceptionally responsive to DAP (RT=3.33), identifying this as a potentially hyper-responsive therapeutic target. However, a UGA at position G542X is predicted to be less responsive to DAP (RT=1.99). Similarly, the model proposes entirely different strategies depending on the specific nucleotide change at a single site; for W1282X, it recommends a DAP-centric approach for a UGA codon but pivots to an eRF1/eRF3 inhibitor like SJ6986 for a UAG codon.

## Discussion

In this study, we developed TransStop, a pan-drug transformer model with a cross-attention mechanism that sets a new standard for predicting drug-induced translational readthrough of PTCs. Our model achieves an overall R^2^ of 0.94 on a held-out test set, significantly increases the predictive power of previous linear models and demonstrates the critical importance of capturing non-linear, context-dependent interactions between nucleotide sequences and therapeutic compounds.

The architectural shift from simple concatenation to cross-attention is central to this success. This mechanism, where a learned drug embedding actively queries the sequence embeddings, allows the model to dynamically assess which parts of the sequence are most relevant for a given drug. The result is a more nuanced and biologically meaningful representation of the drug-sequence interaction. This is demonstrated by the model’s ability to not only predict readthrough efficiency with high accuracy but also to internally organize its latent space according to fundamental biological principles, such as clustering sequences by stop codon type and arranging them along a continuous gradient of readthrough potential, all without explicit supervision.

The practical implications of this advanced model are far-reaching. Our genome-wide analyses revealed clear drug specializations, providing a strong rationale for tailoring nonsense suppression therapy to the specific mutation of a patient. For example, our results establish Clitocine, DAP, and SJ6986 as the best specialists for UAA, UGA, and UAG codons, respectively.

Perhaps most significantly, our work demonstrates that a more powerful model not only yields incremental improvements but also generates qualitatively new and actionable insights. The systematic disagreements with the Toledano et al. model, particularly in the context of UGA codons, highlights the limitations of linear assumptions. TransStop revises therapeutic recommendations in 41,664 cases with a significant predicted gain in efficiency, providing a clear path toward rescuing previously suboptimal treatment strategies. The ability to generate a detailed “therapeutic matrix” for any gene, as demonstrated with CFTR, transforms our predictive tool into a hypothesis-generation engine for precision medicine, enabling rational patient stratification and clinical trial design.

While our model marks a significant advance, it is based on data from a single reporter system and cell line (HEK293T). Future work will aim to integrate data from diverse cellular contexts and incorporate additional features like mRNA secondary structure. Nonetheless, TransStop provides a robust and extensible framework, paving the way for a new generation of predictive tools to accelerate the development of personalized therapies for genetic diseases.

### Perspectives

The development of the TransStop model should make a significant contribution to the advancement of personalized nonsense suppression therapies. By providing a highly accurate and interpretable tool for predicting drug efficacy, TransStop opens the way to a new paradigm in which clinical trial design and patient treatment are guided by robust, data-driven predictions. We envision a future where the genetic sequence of a patient’s PTC can be used to instantly predict the most effective compound from a panel of available drugs, maximizing the potential for therapeutic benefit and minimizing the risk of ineffective treatment.

Furthermore, the framework presented here is readily extensible. As new readthrough-inducing compounds are discovered, they can be integrated into our pan-drug model, continually expanding its predictive power and clinical utility. By combining the power of deep learning with large-scale experimental data, we are moving closer to the ultimate goal of precision medicine: delivering the right drug to the right patient at the right time.

## Acknowledgements

We thank the members of the BiGEst-ICube platform for their assistance.

## Funding Information

This work was supported by institute funds from the French Centre National de la Recherche Scientifique, and the University of Strasbourg.

## Data availability

The source code is available at https://github.com/Dichopsis/TransStop. The TransStop model is available at https://huggingface.co/Dichopsis/TransStop. The genome-wide predictions are available at https://doi.org/10.5281/zenodo.16918476.

## Notes

### Competing Interest Statement

The authors have declared no competing interest.

